# Coronary Microvascular Dysfunction is Associated with Augmented Lysosomal Signaling in Hypercholesterolemic Mice

**DOI:** 10.1101/2024.07.10.603000

**Authors:** Yun-Ting Wang, Alexandra K. Moura, Rui Zuo, Wei Zhou, Zhengchao Wang, Kiana Roudbari, Jenny Z. Hu, Pin-Lan Li, Yang Zhang, Xiang Li

## Abstract

Accumulating evidence indicates that coronary microvascular dysfunction (CMD) caused by hypercholesterolemia can lead to myocardial ischemia, with or without obstructive atherosclerotic coronary artery disease (CAD). However, the molecular pathways associated with compromised coronary microvascular function prior to the development of myocardial ischemic injury remain poorly defined. In this study, we investigated the effects of hypercholesterolemia on the function and integrity of the coronary microcirculation in mice and the underlying mechanisms. Mice were fed with a hypercholesterolemic Paigen’s diet (PD) for 8 weeks. Echocardiography data showed that PD caused CMD, characterized by significant reductions in coronary blood flow and coronary flow reserve (CFR), but did not affect cardiac remodeling or dysfunction. Immunofluorescence studies revealed that PD-induced CMD was associated with activation of coronary arterioles inflammation and increased myocardial inflammatory cell infiltration. These pathological changes occurred in parallel with the upregulation of lysosomal signaling pathways in endothelial cells (ECs). Treating hypercholesterolemic mice with the cholesterol-lowering drug ezetimibe significantly ameliorated PD-induced adverse effects, including hypercholesterolemia, steatohepatitis, reduced CFR, coronary EC inflammation, and myocardial inflammatory cell infiltration. In cultured mouse cardiac endothelial cells (MCECs), 7-ketocholesterol (7K) increased mitochondrial reactive oxygen species (ROS) and inflammatory responses. Meanwhile, 7K induced the activation of TFEB and lysosomal signaling in MCECs, whereas the lysosome inhibitor bafilomycin A1 blocked 7K-induced TFEB activation and exacerbated 7K-induced inflammation and cell death. Interestingly, ezetimibe synergistically enhanced 7K-induced TFEB activation and attenuated 7K-induced mitochondrial ROS and inflammatory responses in MCECs. These results suggest that CMD can develop and precede detectable cardiac functional or structural changes in the setting of hypercholesterolemia, and that upregulation of TFEB-mediated lysosomal signaling in ECs plays a protective role against CMD.

## Introduction

Coronary artery disease (CAD) is the most prevalent form of heart disease and a leading cause of death in the United States and other developed countries ^1^. The coronary arterial system consists of distinct compartments, including large epicardial coronary arteries, arterioles, and capillaries ^2^. The network of coronary arterioles and capillaries, known as the coronary microcirculation, plays a vital role in ensuring normal myocardial perfusion ^2^. Functional or structural abnormalities in the coronary microcirculation can lead to impaired myocardial perfusion, resulting in a condition called coronary microvascular dysfunction (CMD), which can lead to myocardial ischemia ^2–4^. CMD has a higher prevalence in individuals with suspected signs and symptoms of ischemia but no obstructive coronary artery disease (INOCA), particularly in women ^5^. CMD is associated with adverse outcomes and has important prognostic significance for various cardiovascular diseases, especially myocardial ischemia ^6–9^. Coronary flow reserve (CFR) is often used as a clinical indicator to assess CMD in INOCA ^6,10–14^. To date, there is no specific treatment to prevent CMD, and therefore it is imperative to elucidate the molecular pathways associated with CMD.

The mechanisms leading to CMD are diverse and complex, and are understudied compared with large coronary arteries. Coronary microvascular endothelial dysfunction is considered as an important mechanism in the initiation and progression of CMD ^15–17^. Endothelial cells (ECs) play a crucial role in regulating vascular tone by producing and releasing vasoactive relaxing or constricting factors. Endothelium-derived relaxing factors include nitric oxide (NO), vasodilatory prostaglandins, and endothelium-derived hyperpolarization factors ^18,19^. Hydrogen peroxide is considered as one of important hyperpolarization factors in various vascular beds, including human coronary arteries ^20–22^. ECs also produce vasoconstricting factors such as endothelin. In this regard, arterial endothelial dysfunction is often characterized by either an impaired vasodilatory response or an overactive vasoconstrictor response. Mechanistically, endothelial dysfunction is accompanied by increased oxidative stress, elevated production of reactive oxygen species (ROS) and vasoconstrictors, and a gradual decline in NO bioavailability ^23^. In addition to impaired vasomotor responses, endothelial dysfunction involves a shift from a quiescent state to an activated, pro-inflammatory, and pro-thrombotic state, resulting in increased expression of inflammatory mediators and enhanced interaction with platelets and leukocytes ^24^. Moreover, persistent and excessive endothelial dysfunction may lead to structural changes in coronary microcirculation including arterioles wall remodeling and microvascular rarefaction ^25^. These CMD-related pathological changes lead to abnormal myocardial hemodynamics and promote the development of myocardial ischemic injury.

Metabolic disorders, such as hypercholesterolemia, are recognized as important predisposing risk factors for CAD, with or without atherosclerotic obstruction and myocardial infarction ^26–28^. Indeed, hypercholesterolemia is common in patients with INOCA and is even higher in patients with obstructive CAD ^29^. However, there are currently no studies using mouse models to characterize CMD and its associated pathological changes caused by hypercholesterolemia. The lack of mechanistic studies in this field has hampered the development of specific therapeutic interventions for CMD. The aim of this study was to determine the early effects of hypercholesterolemia on the coronary microcirculation and myocardial tissue in mice before the onset of cardiac remodeling and dysfunction. We used a murine hypercholesterolemia model established by high-fat and high-cholesterol Paigen’s diet (PD) feeding, and CMD was characterized by reductions in coronary blood flow and CFR. We evaluated a series of cardiovascular-related pathological changes in the coronary arterioles, capillaries and myocardium, including arterioles wall thickening and lipid accumulation, microvascular rarefaction, thrombosis, and inflammation. We also investigated whether lysosomal signaling was changed in coronary ECs. In addition, we investigated the protective effects of the cholesterol-lowering drug ezetimibe on CMD in hypercholesterolemic mice. We also confirmed the direct effects of ezetimibe on the activation of TFEB-lysosomal signaling and attenuation of EC inflammation and injury in a 7-ketocholesterol (7K)–induced cellular model. Our findings provide new insights that activation of lysosome signaling may be a potential therapeutic target for treating CMD in metabolic disorders.

## Materials and Methods

### Mice

All experimental protocols were reviewed and approved by the University of Houston Institutional Animal Care and Use Committee. Female wild-type mice (B6.129 background) aged 6-12 months were used for all experiments. Mice were housed in a temperature-controlled room with 12-hour dark-light cycle and provided standard rodent chow and water ad libitum. Mice were randomly assigned into a normal chow diet (ND) group or a Paigen’s diet (PD) (Research diet, D12336) group for 8 weeks. For experiments with ezetimibe administration, mice were divided into three groups: ND vehicle control, PD vehicle control, and PD with ezetimibe treatment. Ezetimibe (3 mg/kg) was administrated intraperitoneally every two days. Vehicle control groups received DMSO+PBS injection. Ezetimibe was dissolved in DMSO and diluted with PBS to keep DMSO concentration below 10% of the total injection volume. After 8 weeks, mice were sacrificed, and blood, heart, liver, and aorta samples were collected and stored at - 80°C for further analysis.

### Echocardiography

Transthoracic echocardiography was performed using a Vevo 3100 micro-ultrasound imaging system with an MX550D-0073 probe (VisualSonics Inc., Canada). Mice were anesthetized with 1.5–2% isoflurane (ISO) mixed with 100% medical oxygen and maintained at 1-1.5% isoflurane to control the heart rate between 400 and 500 beats per minute. Heart rate and electrocardiogram were monitored by limb electrodes. Body temperature was monitored and maintained between 36°C and 37°C. M-mode images were obtained from the parasternal short-axis view at the level of the papillary muscles and analyzed with Vevo LAB 2.1.0. Left anterior descending coronary artery flow velocity was measured using pulsed-wave doppler under baseline and hyperemic conditions induced by inhalation of 1.0% and 2.5% isoflurane, respectively. CFR was calculated as the ratio of peak blood flow velocity during hyperemia to baseline.

### Antibodies and reagents

Primary antibodies: CD41 (BD 553847), FITC-αSMA (Sigma F3777), CD45 (Abcam ab25386), F4/80 (BD 565411), Von Willebrand Factor (VWF) (Abcam ab11713), CD41 (BD 553847), NG-2 (Abcam ab275024), ZO-1 (Thermo 617300), caspase-1 (Adipogen AG-20B-0044-C100), HMGB-1 (Abcam ab79823), VCAM-1 (Abcam ab134047), Cathepsin B (Abcam ab58802), LAMP-1 (BD 553792), LAMP-2A (abcam ab18528), TFEB (Bethyl Laboratories A303-673A), TRPML-1 (Thermo PA1-46474), PLIN2 (Proteintech 15294-1-AP), LC3A/B (CST 12741S), CCL-2 (Millipore MABN712), β-actin (CST 3700S).

Secondary antibody for western blot: IRDye® 800CW anti-Mouse IgG (LICOR 926-32212), IRDye® 800CW anti-Rabbit IgG (LICOR 926-32213), anti-Mouse IgG, HRP (Thermo Fisher A16011), stabilized peroxidase conjugated anti-rabbit (Invitrogen 32460), anti-rat IgG-HRP (Fisher 629520). Secondary antibody for immunofluorescence: anti-Mouse IgG, Alexa Fluor 488 conjugate (Thermo Fisher A-21202), Alexa Fluor 488 conjugate anti-Rabbit IgG (Thermo Fisher A21206), Alexa Fluor 555 conjugate anti-Mouse IgG (Thermo Fisher A-31570), Alexa Fluor 555 conjugate anti-Rabbit IgG (Thermo Fisher A-31572).

Reagents: Oil red (VWR BT135140-100G), Isolectin GS-IB4 (Fisher I21411), Enzychrom AF cholesterol Assay Kit (BioAssay E2CH-100), EnzyChrom Triglyceride Assay Kit (BioAssay ETGA-200), Bafilomycin A1 from streptomyces griseus (Sigma B1793), 7-Ketocholesterol (7-keto, Sigma C2394), Ezetimibe (Cayman 16331), mitoSOX® Red mitochondrial superoxide indicator (moleculae probes M36008), Green FLICA Caspase-1 Assay Kit (Immunochemistry Technologies, #98), CCK8 kit (APExBIO, K1018), Aurum Total RNA Mini Kits (Bio-Rad, 732-6820), iScript Reverse Transcription Supermix for RT-qPCR (Bio-Rad, 1708841), iTaq Universal SYBR Green supermix (Bio-Rad, 1725121).

### Immunofluorescence staining

Immunofluorescence staining was performed using frozen tissues or cultured MCECs. For tissue staining, frozen section slides were fixed with 4% paraformaldehyde (PFA) for 15 minutes at room temperature. For MCECs, approximately 7.5×10^4^ cells were seeded into 24-well culture plates containing gelatin-coated coverslips. After treatment with indicated stimuli, MCECs were fixed with 4% PFA for 15 minutes at room temperature. After washing with PBS, frozen tissues or cells were blocked and permeabilized with 5% BSA+ 0.3% Triton X-100 in PBS for 1 hour at room temperature and then incubated with primary antibodies overnight at 4°C. After washing with PBST (PBS+0.05% Tween-20), the slides were incubated with corresponding secondary antibodies conjugated to Alexa Fluor 488 or Alexa Fluor 555 (Invitrogen) for 1 hour at room temperature. After washing with PBST, slides were mounted by DAPI-mounting solution and then analyzed by using Olympus IX73 imaging system. The Pearson’s correlation for co-localization efficiency or mean fluorescence density was analyzed using the Image-Pro Plus 6.0 software, as described previously ^30^.

### Hematoxylin and eosin (H&E) staining

H&E staining was performed using hematoxylin and eosin staining kit (Teomics HAE-1). Heart or liver frozen section slides were fixed with 4% PFA for 15 minutes at room temperature. After washing with distilled water, tissues were incubated with adequate Hematoxylin, Mayer’s for 5mins, followed with 2 changes of distilled water to remove excess stain. Then bluing reagent was added to tissues for 10-15 seconds followed with 2 changes of distilled water. After dipping slides in absolute alcohol and wiping excess off, tissues were incubated with adequate Eosin Y solution for 2-3 minutes, followed with rinse using absolute alcohol. Then clear the slide and mount in synthetic resin. Image using Olympus IX73 imaging system promptly.

### Oil Red O staining

Oil Red O stock solution: saturated oil red O in isopropanol (0.3g oil red O in 100 mL isopropanol). Using gentle heat from a water bath to dissolve. Oil red O working solution (fresh): 30 mL of stock solution is mixed with 20 mL distilled water, let stand for 10 minutes. Filter twice with a syringe filter, and use the working solution within 1 hour.

Staining procedure: Heart or liver frozen sections were prepared at a thickness of 8 µm. Air dried sections onto slides, fixed in 4% PFA for 15min, and rinsed immediately in 3 changes of distilled water before being soaked for 20 min. The sections were then washed with 60% isopropanol for 1min to avoid water carryover. Sections were stained with freshly prepared Oil Red O working solution for 30 min, and then rinsed with 60% isopropanol for 2min followed by 2 changes of distilled water. Nuclei were lightly stained with alum hematoxylin for 1min, then rinsed in running tap water for 10min. Sections were mounted in mounting media and images were taken promptly with the Olympus IX73 imaging system.

### Quantitative Real-time PCR

Aurum Total RNA Mini Kits (Bio-Rad, 732-6820) were used to isolate total RNA from heart, aorta, and MCECs. iScript Reverse Transcription Supermix (Bio-Rad, 1708841) was used to generate cDNA from isolated total RNA. The Real-Time PCR was performed using the iTaq Universal SYBR Green supermix (Bio-Rad, 1725121) on the Bio-Rad CFX Connect Real-Time System. Primers of NLRP3, NLRP-1A, ASC, Caspase-1, IL-1beta, IL-18, IL-6, IL-8, GSDMD, TNF, ICAM-1, VCAM-1, P-selectin, E-selectin were purchased from Bio-Rad. Other primers used in the present study were listed in the supplementary materials. The cycle threshold values were converted to relative gene expression levels using the 2-ΔΔCt method. The data was normalized to internal control β-actin or GAPDH ^31^.

### Total cholesterol and triglyceride in serum or liver homogenate

Total cholesterol and triglyceride levels in serum or liver homogenate were measured using a cholesterol assay kit (BioAssay E2CH-100) and a triglyceride assay kit (BioAssay ETGA-200), respectively. Briefly, for the cholesterol assay, 55 uL of assay buffer was mixed with 1 uL of enzyme mix and 1 uL of dye reagent before 50 uL of this working reagent was added to diluted 50 uL standard and sample wells. For the triglyceride assay, working reagent was prepared by mixing 100 uL assay buffer, 2 uL enzyme mix, 5 uL lipase, 1uL ATP, and 1 uL dye reagent in each well. 100 uL of this working reagent mix was added to each diluted 10 uL standard and samples well. The plates were tapped to mix and incubated at room temperature for 30 minutes before having the OD measured at 570 nM with a microplate reader (BMG Labtech). The cholesterol and triglyceride concentrations in the liver homogenate were normalized to protein concentration.

### Mouse cardiac endothelial cell culture

Mouse cardiac endothelial cells (MCECs, Cedarlane, CLU510) were cultured in low glucose DMEM with 5% FBS, 1% penicillin/streptomycin, and 1mM HEPEs at 37°C with 5% CO_2_.

### Western blotting

MCECs were lysed in Laemmli sample buffer (Bio-Rad, 161-0737) containing β-mercaptoethanol (Sigma Aldrich, M3148) and boiled for 10 min at 95 °C before being placed in an ice cooled ultrasonic bath for 5 min. The prepared samples were separated by 12% sodium dodecyl sulfate-polyacrylamide gel electrophoresis. The proteins from these samples were then electrophoretically transferred onto a PVDF membrane at 35 V at 4 °C overnight. The membranes were blocked with 5% BSA in Tris-buffered saline with 0.05% Tween 20. After further washing, the membranes were incubated with primary antibodies as indicated according to the manufacturer’s instructions. After washing with PBST, the membranes were then incubated with corresponding secondary antibodies for 1 hour at room temperature. Finally, the bands were washed with PBST, visualized and analyzed by the LI-COR® Odyssey Fc System.

### Mitochondrial ROS

Mitochondrial ROS production was measured using the MitoSOX™ Red mitochondrial superoxide indicator (molecular probes M36008). Approximately 7.5×10^4^ MCECs were seeded on coverslips in 24-well plates in DMEM with 5% FBS. After treatments, cells were incubated with 5µM MitoSOX™ in culture medium for 10 minutes at 37 °C in the dark. Cells were washed with warmed PBS three times and incubated with Hoechst 33342 (0.5% v/v) for 5 minutes to stain nuclei. Coverslips were then mounted on slides for examination. Images were taken immediately using the Olympus IX73 imaging system.

### Caspase-1 activity assay by FAM-FLICA kit

Approximately 7.5×10^4^ MCECs were seeded on coverslips in 24-well plates for overnight, and the medium was replaced with DMEM containing 1% FBS. Cells were pretreated with or without 50 nM bafilomycin A1 for 1 hour, then co-treated with or without 40 µM 7K for 24 hours. Subsequently, 1x FAM-FLICA (diluted by culture media) was added to the cells on the coverslips for 1 hour. The medium on the coverslips was carefully removed and replaced with apoptosis wash buffer for 10 minutes. Wash buffer was then removed, and cells were stained with Hoechst 33342 at 0.5% v/v for 5 minutes in order to stain the nucleus. To identify dead cells, cells were incubated with propidium iodide (PI, 0.5% v/v) for 5 minutes. Cells were then fixed with a fixative solution at a v/v ratio of 1:10 in PBS for 15 minutes at room temperature. After fixation, coverslips were mounted on slides for examination. Images were taken immediately using the Olympus IX73 imaging system.

### Cell viability assay

In brief, 1.0×10^4^ MCECs were seeded into 96-well plates overnight, and then the medium was replaced with DMEM containing 1% FBS. Cells were treated as described, and 10 µl CCK8 solution was added to each well and incubated at 37 °C for 1 hour. The absorbance was measured at 450 nm using a microplate reader (BMG Labtech).

### Calcein-AM labeled monocytes retention

Macrophage J774 cells were trypsinized and stained with 2 µM fluorescent Calcein-AM (Thermo Fisher, C3100MP) at 37 °C for 10 minutes. After treatment as indicated, 5 x 10^4^ Calcein-AM labeled J774 cells were seeded onto MCECs for 10-30 minutes. Unbounded monocytes were discarded, and MCECs layers with attached monocytes were gently washed with PBS, followed by fixation with 4%PFA for 10 minutes at room temperature. The nucleus was stained with DAPI for 15 minutes at room temperature and mounted with an anti-fluorescence quenching agent. Imaging was performed using the Olympus IX73 imaging system.

### Statistics analysis

Data are presented as Mean±Std. Error of Mean. All experiments were analyzed by the one or two-way ANOVA with different treatments as category factors, followed by Bonferroni’s multiple comparisons test if applicable. A Student’s t-test was used to detect significant difference between two groups. Statistical analysis was carried out using GraphPad Prism 6.0 software (GraphPad Software, USA). P<0.05 was considered statistically significant.

## Results

### Effects of PD on CFR, cardiac remodeling, and heart function

CMD was measured by non-invasive echocardiography, and assessed by coronary blood flow and CFR. Representative images and summarized data (**Fig.1A and 1B**) indicate that PD treatment significantly decreased coronary blood flow velocity in hyperemic mice (2.5 % isoflurane) at 6 and 8 weeks compared with ND group, but had no effects at the baseline (1.0 % isoflurane), corresponding to a decrease in CFR. Next, we evaluated whether PD affected cardiac remodeling (**Fig.1C**) and heart function (**Fig.1D**). We found that PD for 8 weeks showed no significant effect on cardiac remodeling, as measured by diastolic left ventricular anterior wall (LVAW;d), systolic left ventricular anterior wall (LVAW;s), diastolic left ventricular posterior wall (LVPW;d), systolic left ventricular posterior wall (LVPW;s), and calculated left ventricular mass (**Fig.1C**). PD for 8 weeks had no significant effect on heart function, as measured by the diastolic left ventricular internal end (LVID;d), systolic left ventricular internal end (LVID; s), diastolic left ventricular volume (LV Vol; d), systolic left ventricular volume (LV Vol;s), and calculated left ventricular ejection fraction (LVEF) and left ventricular shortening fraction (LVFS) (**Fig.1D**). These data demonstrate that PD causes CMD without affecting cardiac remodeling or heart function.

**Figure 1.**
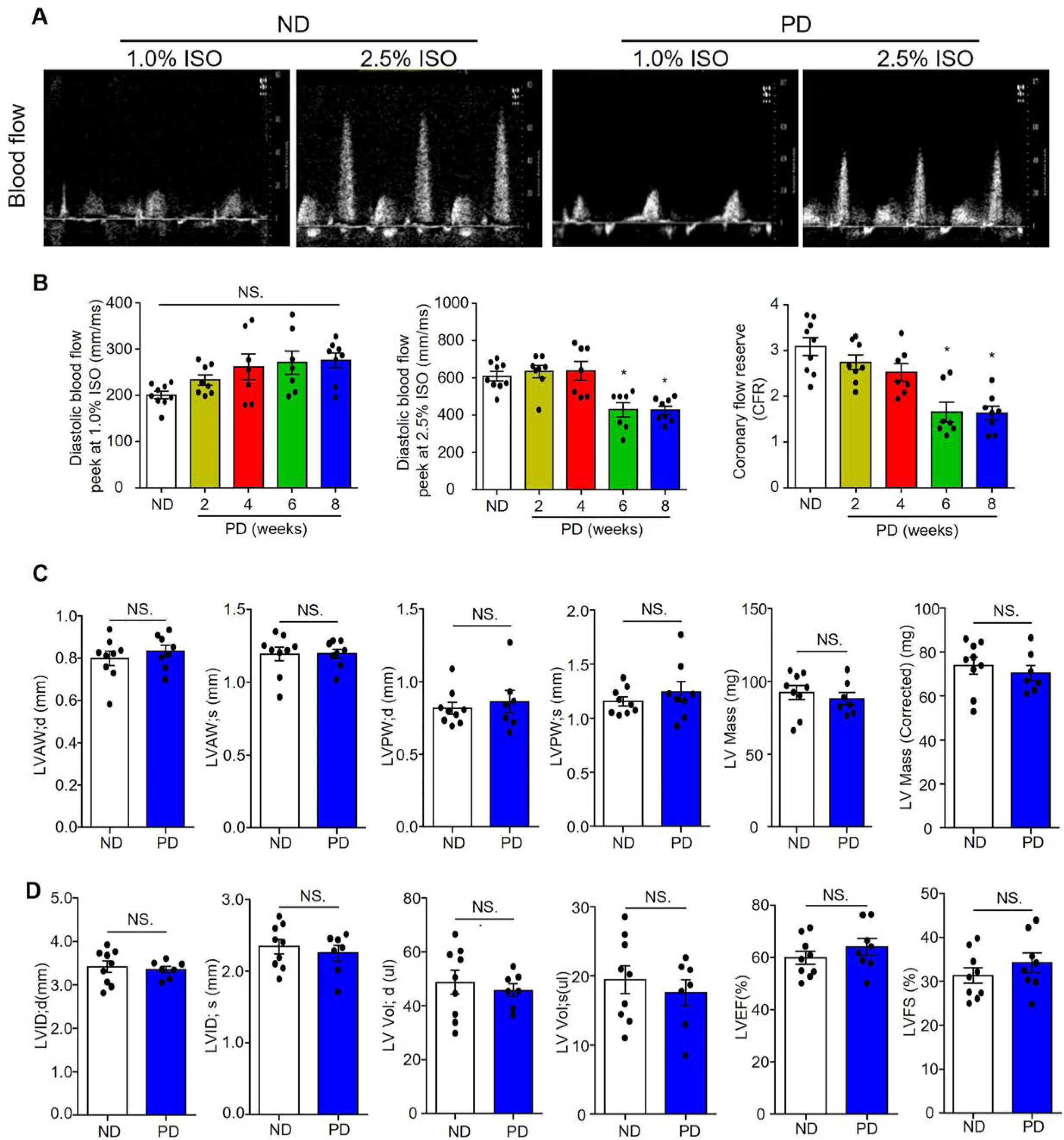
Effects of PD on coronary flow reserve (CFR), cardiac remodeling and heart function. **A,** Representative ultrasound images of isoflurane-induced vasodilation of the left anterior descending coronary artery at basal (1.0% ISO) and hyperemia (2.5% ISO) levels in mice fed with ND or PD for 8 weeks. **B,** Summarized data of diastolic blood flow velocity peeks at 1.0 % ISO, 2.5% ISO, and CFR (ratio of 2.5% ISO over 1.0% ISO) in mice fed with ND or PD for 2, 4, 6, and 8 weeks. **C,** Summarized data of echocardiographic parameters in cardiac remodeling: diastolic left ventricular anterior wall (LVAW; d), systolic left ventricular anterior wall (LVAW; s), diastolic left ventricular posterior wall (LVPW; d), systolic left ventricular posterior wall (LVPW; s), left ventricle mass (LV Mass), corrected LV mass in mice fed with ND or PD for 8 weeks. **D,** Summarized data of echocardiographic parameters in heart function: diastolic left ventricular internal end (LVID; d), systolic left ventricular internal end (LVID; s), diastolic left ventricle volume (LV Vol; d), systolic left ventricle volume (LV Vol; s), left ventricular ejection fraction (LVEF), and left ventricular fractional shortening (LVFS) in mice fed with ND or PD for 8 weeks. \**P*< 0.05, NS. No Significance, (n=7-9).

### Effects of PD on coronary arterioles remodeling, capillary rarefaction, atherothrombosis events, and arterioles and cardiac inflammation

Next, we characterized PD-induced CMD by examining various cardiovascular-related pathological changes. As shown in **Fig.2A**, H&E staining showed that no neointima formation or changes in media thickness were observed in coronary arterioles in both ND and PD groups. These results indicate that PD for 8 weeks did not induce coronary arterioles remodeling. We also performed immunofluorescence analysis of myocardial capillary density by staining microvascular ECs with isolectin-IB4 and pericytes with NG2. As shown in **Fig.2B**, PD did not affect IB4 or NG2 staining, nor did it alter the ratio of NG2 to IB4, suggesting that PD does not alter capillary density or pericyte coverage. Interestingly, Oil Red O staining in **Fig.2C** showed that lipid deposition/fatty streaks were detected in the coronary arterioles walls in 3 of 8 PD-fed mice but not in the ND group. As shown in **Fig.2D**, immunofluorescence analysis detected CD41-positive thrombus in some coronary arterioles in 2 of 8 PD-fed mice, but not in the ND group. Furthermore, PD increased inflammasome activation in the endothelium of coronary arterioles, as evidenced by the increased co-localization (yellow color) of caspase-1 **(Fig.2E)** or high mobility group box 1 (HMGB-1) (**Fig.2F**) with von Willebrand factor (VWF) (EC marker). PD also increased the expression of vascular cell adhesion molecule 1 (VCAM-1) in the endothelium of arterioles **(Fig.2G)**, and increased the adhesion of CD45-positive leukocytes in cardiac tissue **(Fig.2H)**. These results suggest that PD-induced CMD is associated with increased activation of coronary arterioles inflammation and thrombosis and increased myocardial inflammatory cell infiltration, but not with coronary arterioles remodeling or capillary rarefaction.

**Figure 2.**
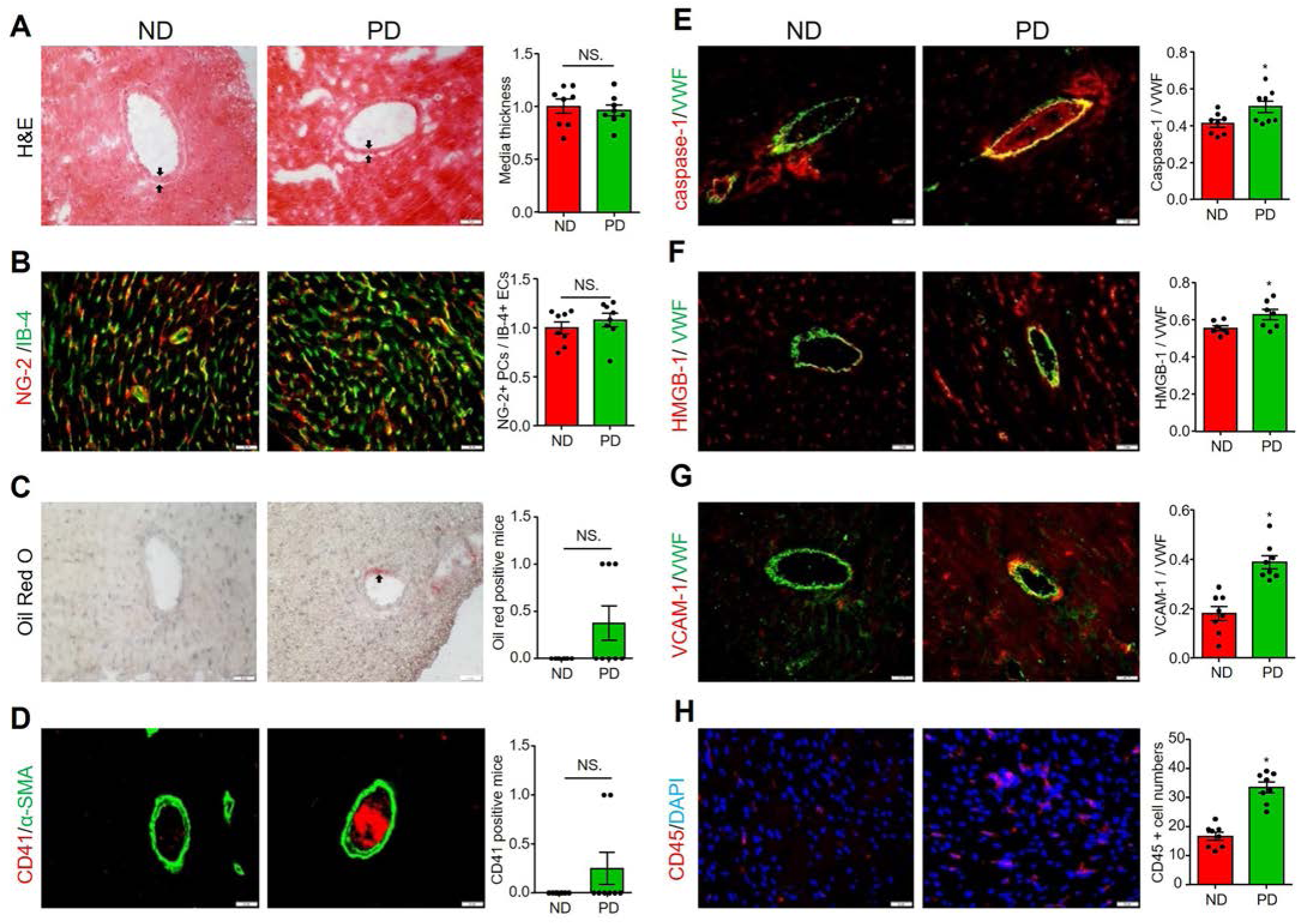
Effects of PD on coronary vascular remodeling, capillary rarefaction, atherothrombosis events, and vascular and cardiac inflammation. All heart samples were collected from mice fed with ND or PD for 8weeks. **A,** Representative H&E staining and summary of coronary artery media thickness (black arrow). **B,** Representative capillary endothelial cell marker Isolectin-IB-4 staining plus pericyte marker NG-2 staining, and summary of colocalization coefficient. **C,** Representative oil Red O staining for lipid deposition/fatty streaks (black arrow) in the coronary artery, and summary of oil Red O staining positive mice numbers. **D,** Representative CD41 staining for platelet activation, and summary of positive mice. Alpha-SMA staining was used to localize coronary arterioles. Representative images for caspase-1/VWF (**E**), HMGB-1/VWF (**F**), and VCAM-1/VWF (**G**) staining and their summarized colocalization coefficients. Von Willebrand factor (VWF) staining was used to localize coronary arterioles endothelium. Cardiac inflammatory cells infiltration was indicated by positive staining of leukocyte marker CD45 **(H)**. Scale bar=20 µm. \**P*< 0.05, NS. No Significance, (n=6-8).

### Effect of PD on transcriptional levels of lipogenesis, thrombosis, and inflammation related pathways

To confirm the pathological changes associated with CMD, we analyzed the expression levels of genes involved in lipid accumulation, thrombosis, and inflammation in the heart or aortas of mice fed with ND or PD for 8 weeks. We first examined genes related to lipid droplet (LD) biogenesis and formation, including glycerol-3-phosphate acyltransferase 4 (GPAT-4), 1-acylglycerol-3-phosphate O-acyltransferase 2 (AGPAT-2), LIPIN-2, diacylglycerol O-acyltransferase 1/2 (DGAT-1, DGAT-2), perilipins (PLIN1, PLIN2, PLIN3, PLIN4, PLIN5), and genes related to cholesterol biosynthesis, including acetyl-CoA acetyltransferase 1/2 (ACAT1, ACAT2). As shown in **Fig.3A**, we found that PD had no significant effects on most of above-mentioned genes in heart tissues except PLIN1, a major perilipin isoform expressed in adipocytes. In contrast, PD significantly increased GPAT-4, AGPAT-2, DGAT-1/2, ACAT1/2, and PLIN5 in aortas. These results suggest that PD treatment tends to increase lipogenesis and LD formation in the arterial wall.

**Figure 3.**
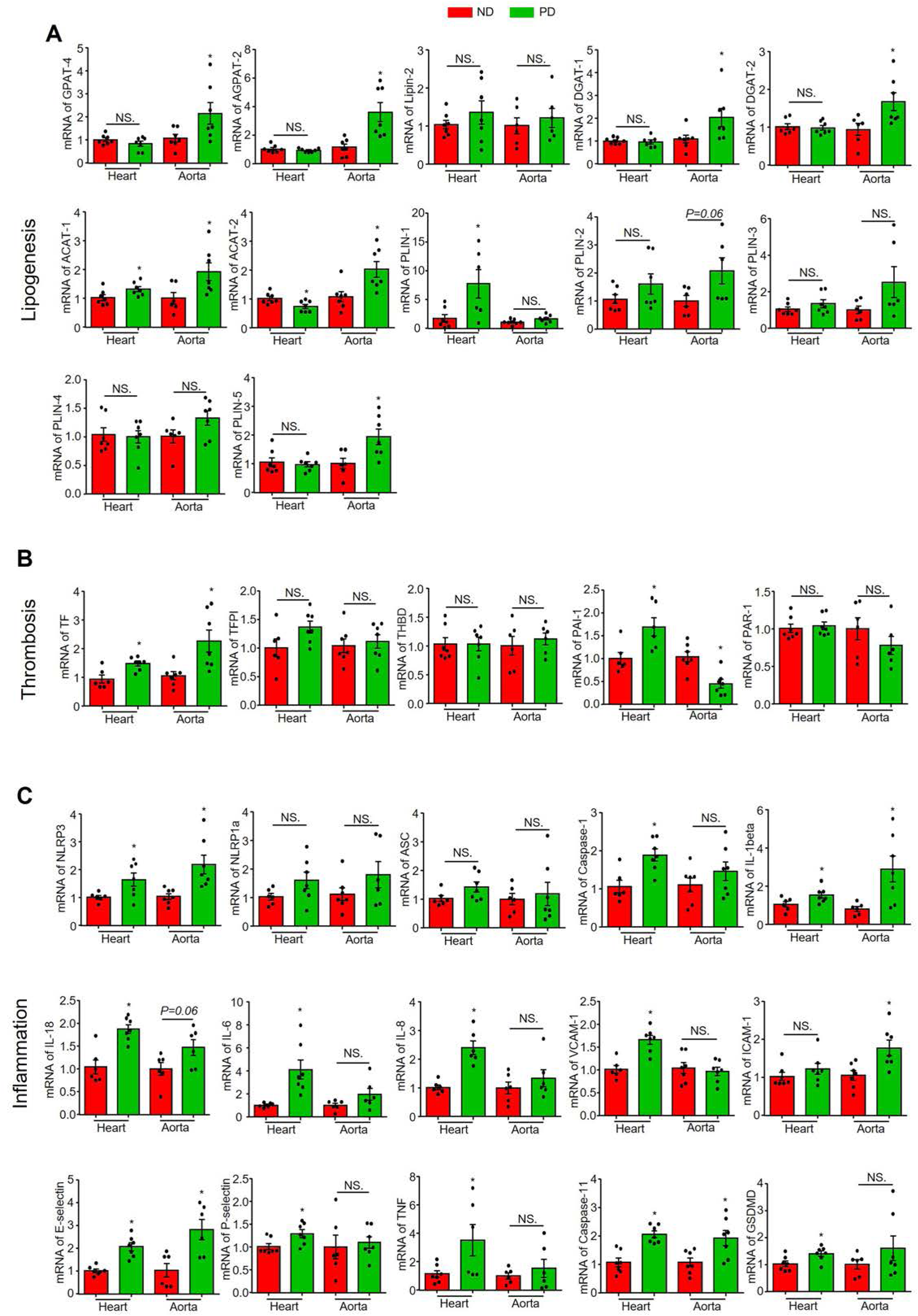
Effect of PD on the transcriptional levels of lipogenesis, thrombosis, and inflammation pathways-related genes. Heart or aorta tissue samples were collected from mice fed with ND or PD for 8weeks. **A,** mRNA levels of lipogenesis related genes: GPAT-4, AGPAT-2, Lipin-2, DGAT-1, DGAT-2, ACAT-1, ACAT-2, PLIN-1, PLIN-2, PLIN-3, PLIN-4, and PLIN-5. **B,** mRNA levels of thrombosis related genes: TF, TFPI, THBD, PAI-,1and PAR-1. **C,** mRNA levels of inflammation related genes: NLRP3, NLRP1a, ASC, caspase-1, IL-1beta, IL-18, IL-6, IL-8, VCAM-1, ICAM-1, E-selectin, P-selectin, TNF, caspase-11, and GSDMD. \**P*< 0.05, NS. No Significance, (n=6-7).

Then, we analyzed thrombosis-related genes including tissue factor (TF), tissue factor pathway inhibitor (TFPI), thrombomodulin (THBD), proteinase-activated receptor 1 (PAR-1), and plasminogen activator inhibitor 1 (PAI-1). As shown in **Fig.3B**, PD increased TF in both hearts and aortas. Additionally, PD increased PAI-1 in the heart but decreased it in the aorta. These results suggest that thrombotic pathways such as coagulation and fibrinolysis are altered in the arterial wall and myocardium.

As for inflammation, we analyzed genes of inflammasomes including NLRP3, NLRP1a, ASC, caspase-1, caspase-11, GSDMD, IL-1β, and IL-18, adhesion molecules including VCAM-1, ICAM-1, E-selectin, and P-selectin, and inflammatory mediators including IL-6, IL-8, and TNFα. As shown in **Fig.3C**, PD significantly upregulated NLRP3, caspase-1, caspase-11, GSDMD, IL-1β, and IL-18 in the heart, but only upregulated NLRP3, IL-1β, and caspase-11 in the aorta. These results suggest the NLRP3 inflammasome is activated in both the arterial wall and the myocardium, with more pronounced effects in the myocardium. PD also upregulated genes for various adhesion molecules, including VCAM-1, E-selectin, and P-selectin in the heart and ICAM-1 and E-selectin in the aorta (**Fig.3C**). Interestingly, PD upregulated inflammatory mediators IL-6, IL-8, and TNFα only in the heart but not in the aorta (**Fig.3C**), which is consistent with the increased adhesion of CD45-positive leukocytes in cardiac tissue.

### PD upregulated lysosomal signaling pathway in coronary arterioles

Recent studies have shown that lysosomal signaling pathways play a critical role in regulating endothelial homeostasis under various metabolic stresses, including hypercholesterolemia ^32,33^. Here, we investigated whether hypercholesterolemia affects the lysosomal signaling in the coronary microcirculation by immunofluorescence studies. The expression of several lysosomal markers was significantly increased in the endothelium of coronary arterioles of mice fed with PD for 8 weeks, including transcription factor EB (TFEB) **(Fig.4A)**, lysosomal-associated membrane protein 1 (LAMP-1) **(Fig.4B)**, lysosomal-associated membrane protein 2A (LAMP-2A) **(Fig.4C)**, and transient receptor potential cation channel, mucolipin subfamily, member 1 (TRPML-1 or Mucolipin-1) **(Fig.4D)**. Furthermore, we quantified the mRNA levels of several lysosome/autophagy-related genes in the heart and aorta tissues. As shown in **Fig.4E**, PD only upregulated the mRNA levels of LAMP-2A and Beclin-1 in the heart, while almost all tested lysosome/autophagy-related genes were significantly upregulated in the aorta, including LAMP-1, TRPML-1, SMPD1, LC3, and p62. Notably, PD did not affect TFEB mRNA levels in aorta and heart tissues (**Fig.4E)**. Nonetheless, these data suggest that PD significantly activates the lysosome signaling pathway in coronary arterioles endothelium, and to a less extent, in the myocardium as well.

**Figure 4.**
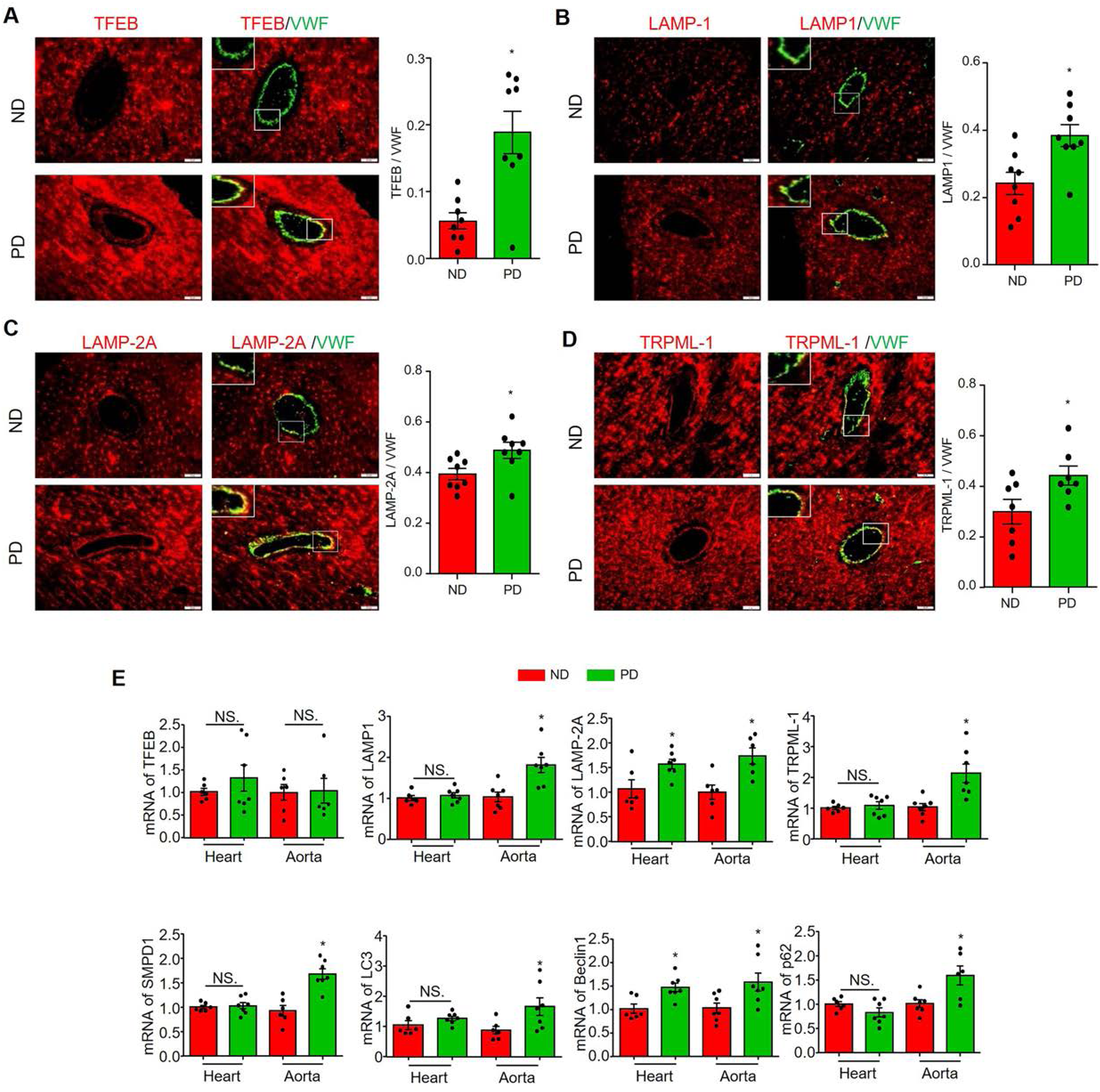
PD upregulated lysosome related signaling pathway in coronary arterioles. Heart or aorta tissue samples were collected from mice fed with ND or PD for 8weeks. Representative images for staining of TFEB/VWF (**A**), LAMP-1/VWF (**B**), LAMP-2A/VWF (**C**), and TRPML-1/VWF (**D**) and the summary of colocalization coefficients. VWF staining was used to localize coronary arterioles endothelium. **E,** mRNA levels of lysosome-autophagy related genes in heart or aorta tissues: TFEB, LAMP-1, LAMP-2A, TRPML-1, SMPD-1, LC3, Beclin-1 and p62. AOI, area of interest, scale bar=20 µm. \**P*< 0.05, NS. No Significance, (n=6-8).

### Ezetimibe alleviated PD-induced steatohepatitis and hypercholesterolemia

Ezetimibe is an FDA-approved drug for the treatment of hypercholesterolemia that works by inhibiting the absorption of dietary and biliary cholesterol from the small intestine. Here, we investigated whether ezetimibe could prevent the development of CMD in mice fed with PD for 8 weeks. We first examined the efficacy of ezetimibe administration by analyzing pathological features of the liver, including steatosis, inflammation, and serum cholesterol levels. As shown in **Fig.5A**, H&E staining showed that ezetimibe prevented steatosis which is confirmed by a decrease in oil red O staining (**Fig.5B and 5E**) and a decrease in the expression of lipid droplet coating protein PLIN2 (**Fig.5C and 5F**). Additionally, ezetimibe attenuated the PD-induced increase in CD45-positive inflammatory cells in the liver (**Fig.5D and 5G**). Next, we measured the total cholesterol and triglyceride levels in the serum and liver tissue. Consistent with the beneficial effects of ezetimibe on steatohepatitis, we found that PD induced an increase in serum and liver cholesterol levels, which were significantly attenuated by ezetimibe administration (**Fig.5H and 5I**). Notably, PD decreased serum triglycerides, but did not affect liver triglycerides, whereas ezetimibe had no further effect (**Fig.5J and 5K**).

**Figure 5.**
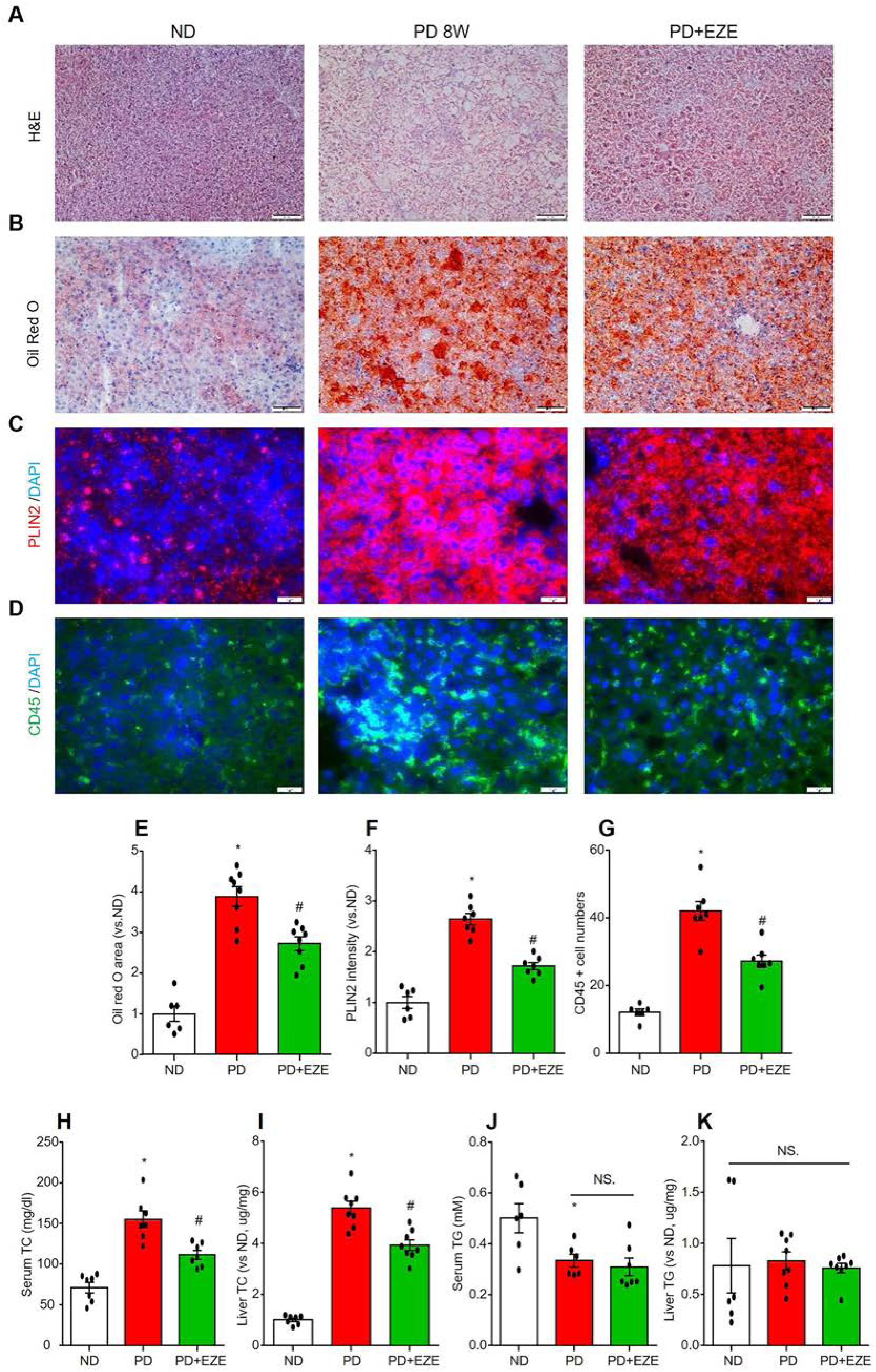
Ezetimibe alleviated PD-induced liver steatohepatitis and hypercholesterolemia. Liver sections, serum, and liver homogenate were collected from mice in the ND-fed vehicle control mice group (PBS+DMSO), the 8-weeks PD-fed vehicle control group (PBS+DMSO) and the PD + ezetimibe (EZE) treatment group (I.P. 3mg/kg/2d for 8 weeks). Liver histomorphology was measured using H&E staining **(A)**. Liver lipid deposition was detected by Oil Red O staining **(B and E)** and lipid droplet associated protein PLIN2 staining **(C and F)**. Liver inflammatory cell infiltration was indicated by leukocyte marker CD45 staining **(D and G)**. Total cholesterol (TC) **(H and I)** and triglyceride (TG) **(J and K)** levels in serum and liver homogenate were measured. Scale bar=20 µm. * vs ND, # vs PD, *P*< 0.05, NS. No Significance, (n=6-8).

### Ezetimibe rescued coronary microvascular function in mice fed with PD

As illustrated in **Fig.6A and 6B**, the administration of ezetimibe to PD-fed mice significantly restored hyperemic coronary blood flow velocity and CFR compared with PD only group. In contrast, ezetimibe had no effect on cardiac remodeling or heart function (**Fig.6C**). As expected, ezetimibe inhibited the PD-induced increase of VCAM-1 (**Fig.7A**) and HMGB-1 (**Fig.7B**) expression in coronary arterioles and capillary ECs. Furthermore, ezetimibe prevented the PD-induced infiltration of CD45-positive inflammatory cells around coronary arterioles and capillaries (**Fig.7C**). These results suggest that ezetimibe prevents PD-induced CMD and inflammatory responses in the coronary microcirculation.

**Figure 6.**
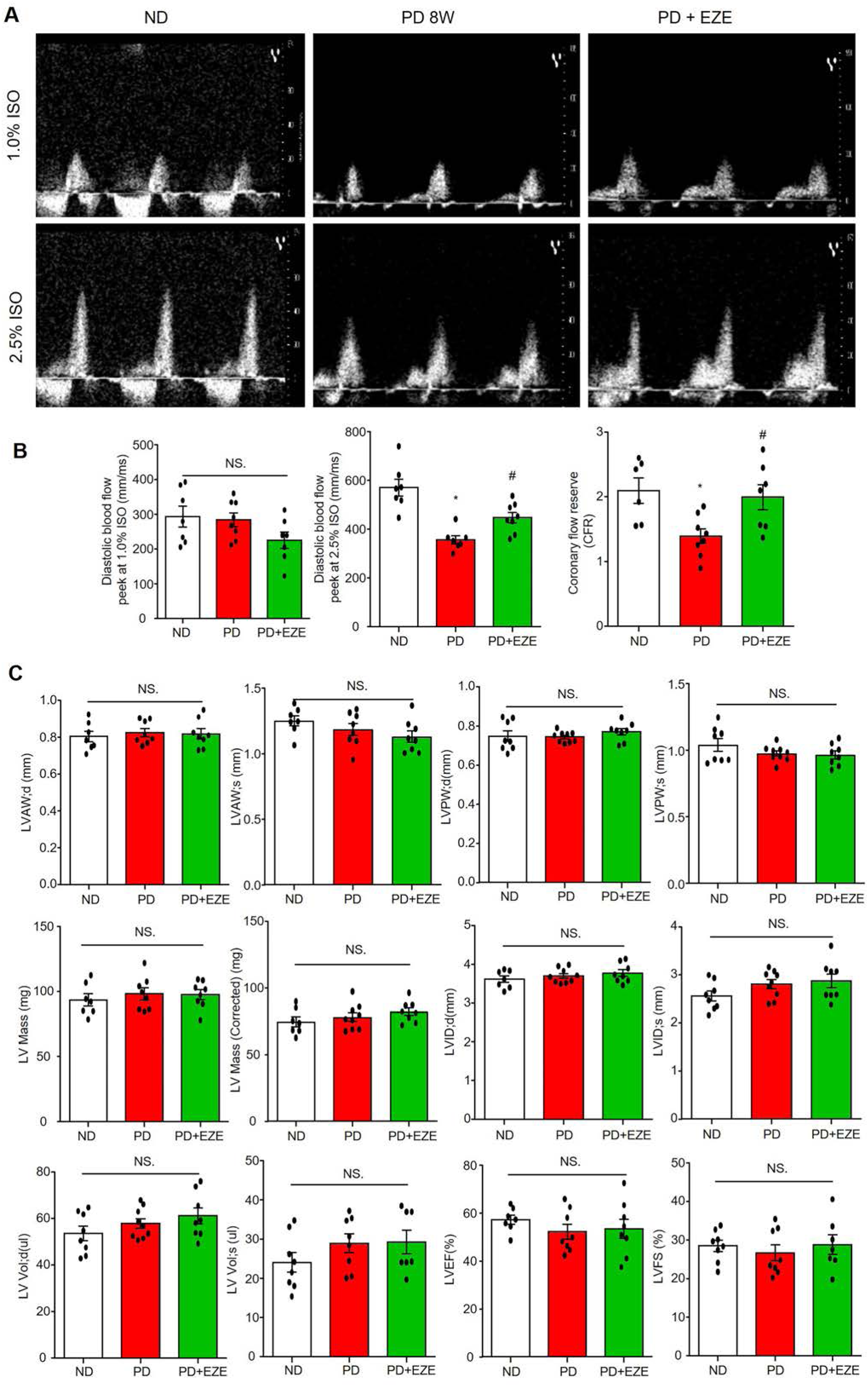
Ezetimibe restored PD-induced CFR decrease. Echocardiographic parameters were measured in mice from the ND-fed vehicle control mice group (PBS+DMSO), the 8-weeks PD-fed vehicle control group (PBS+DMSO) and the PD + ezetimibe (EZE) treatment group (I.P. 3mg/kg/2d for 8 weeks). **A,** Representative ultrasound images of isoflurane-induced vasodilation of the left anterior descending coronary artery at basal (1.0% ISO) and hyperemia (2.5% ISO) levels. **B,** Summarized data of diastolic blood flow velocity peeks at 1.0% ISO, 2.5% ISO and CFR (ratio of 2.5% ISO over 1.0% ISO). **C,** Summarized data of echocardiographic parameters in cardiac remodeling and heart function: (LVAW; d), (LVAW; s), (LVPW; d), (LVPW; s), LV Mass, corrected LV mass, (LVID; d), (LVID; s), (LV Vol; d), (LV Vol; s), LVEF, and LVFS. * vs ND, # vs PD, *P*< 0.05, NS. No Significance, (n=7-9).

**Figure 7.**
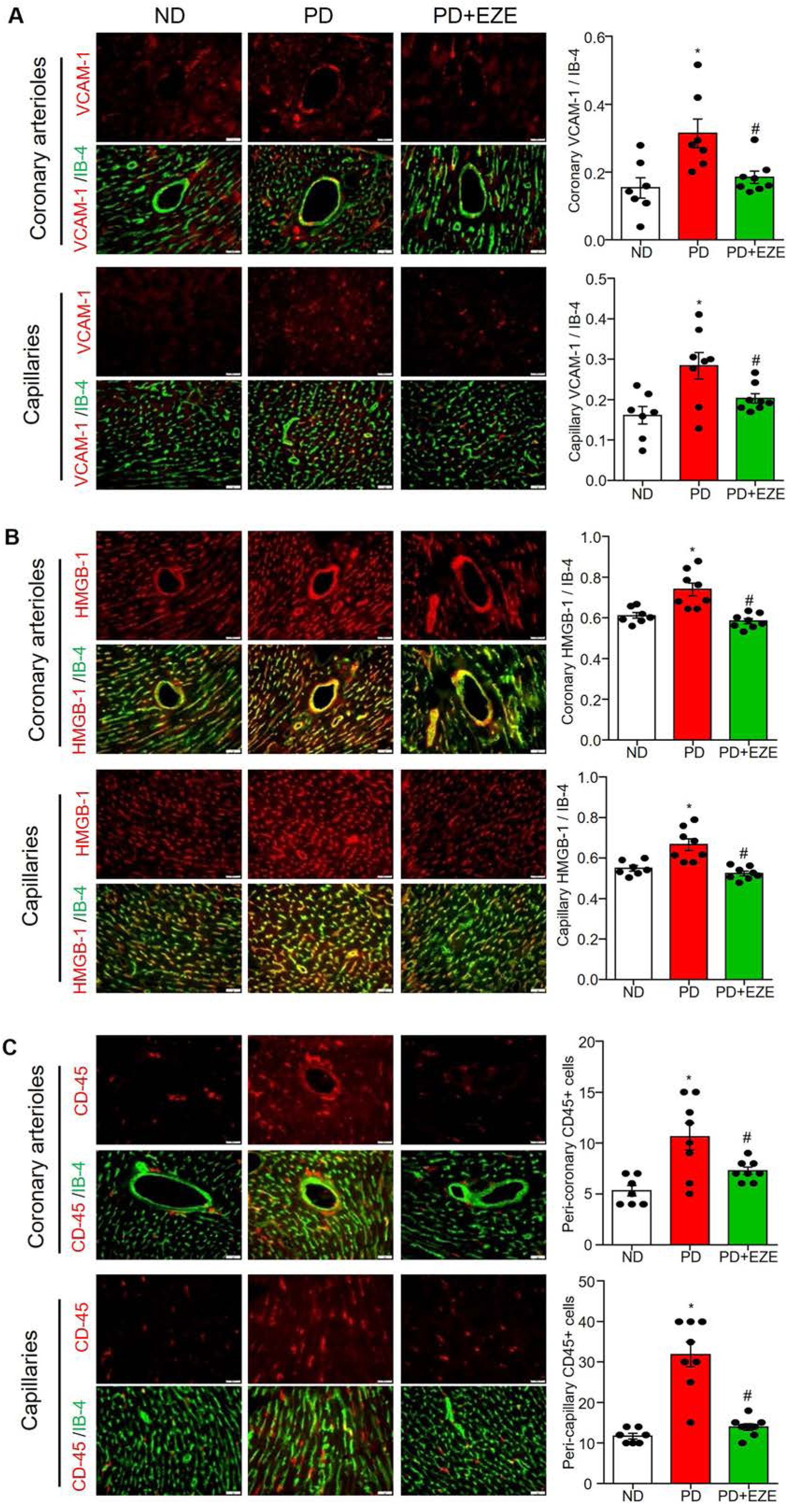
Ezetimibe reduced PD-induced inflammation and monocyte adhesion in both coronary arterioles and capillaries. Heart sections collected were from mice in the ND-fed vehicle control mice group (PBS+DMSO), the 8-weeks PD-fed vehicle control group (PBS+DMSO) and the PD + ezetimibe (EZE) treatment group (I.P. 3mg/kg/2d for 8 weeks). The staining and summarized colocalization coefficients or expression of VCAM-1/IB4 **(A)**, HMGB-1/IB4 **(B)**, and CD45 **(C)** in coronary arterioles endothelium and capillary endothelial cells. Isolectin-IB-4 staining was used to localize coronary arterioles endothelium and capillary endothelial cells. Scale bar=20 µm. * vs ND, # vs PD, *P*< 0.05, NS. No Significance, (n=6-8).

### Mitochondrial ROS production and inflammation is associated with upregulation of lysosomal signaling in cultured MCECs

Next, we investigated the association between adverse effects and augmented lysosomal signaling in cultured MCECs. First, we confirmed that 7-ketocholesterol (7K) dose-dependently increased mitochondrial superoxide production as measured by mitoSOX Red staining (**Fig.8A and 8D**). 7K treatment also significantly upregulated the expression of VCAM-1 (**Fig.8B and 8E**) and C-C motif chemokine ligand 2 (CCL2) (**Fig.8C and 8F**). These results suggest that 7K induces mitochondrial ROS production and inflammation in cultured MCECs, which mirrors the pathological changes in coronary ECs under hypercholesterolemic conditions *in vivo*. We then examined the effects of 7K on the lysosomal signaling pathway by analyzing the activation of transcription factor EB (TFEB), a master regulator of genes involved in autophagy and lysosomal signaling ^31,34^. Once activated, TFEB translocates from the cytoplasm to the nucleus. As shown in **Fig.8G**, 7K dose-dependently increased the nuclear translocation of TFEB in MCECs. Western blot analysis **(Fig.8H)** showed that 7K increased the protein expression of the autophagosome marker LC3A/B-II. Furthermore, 7K significantly increased the mRNA levels of genes in autophagy and lysosomal signaling pathways, including LAMP-2A, Beclin-1, LC3 and p62 **(Fig.8I)**. In contrast, 7K had no effect on the LAMP-1 and even suppressed the mRNA level of TFEB **(Fig.8I)**. These results suggest that 7K activates TFEB-mediated autophagy and lysosomal signaling in cultured MCECs.

**Figure 8.**
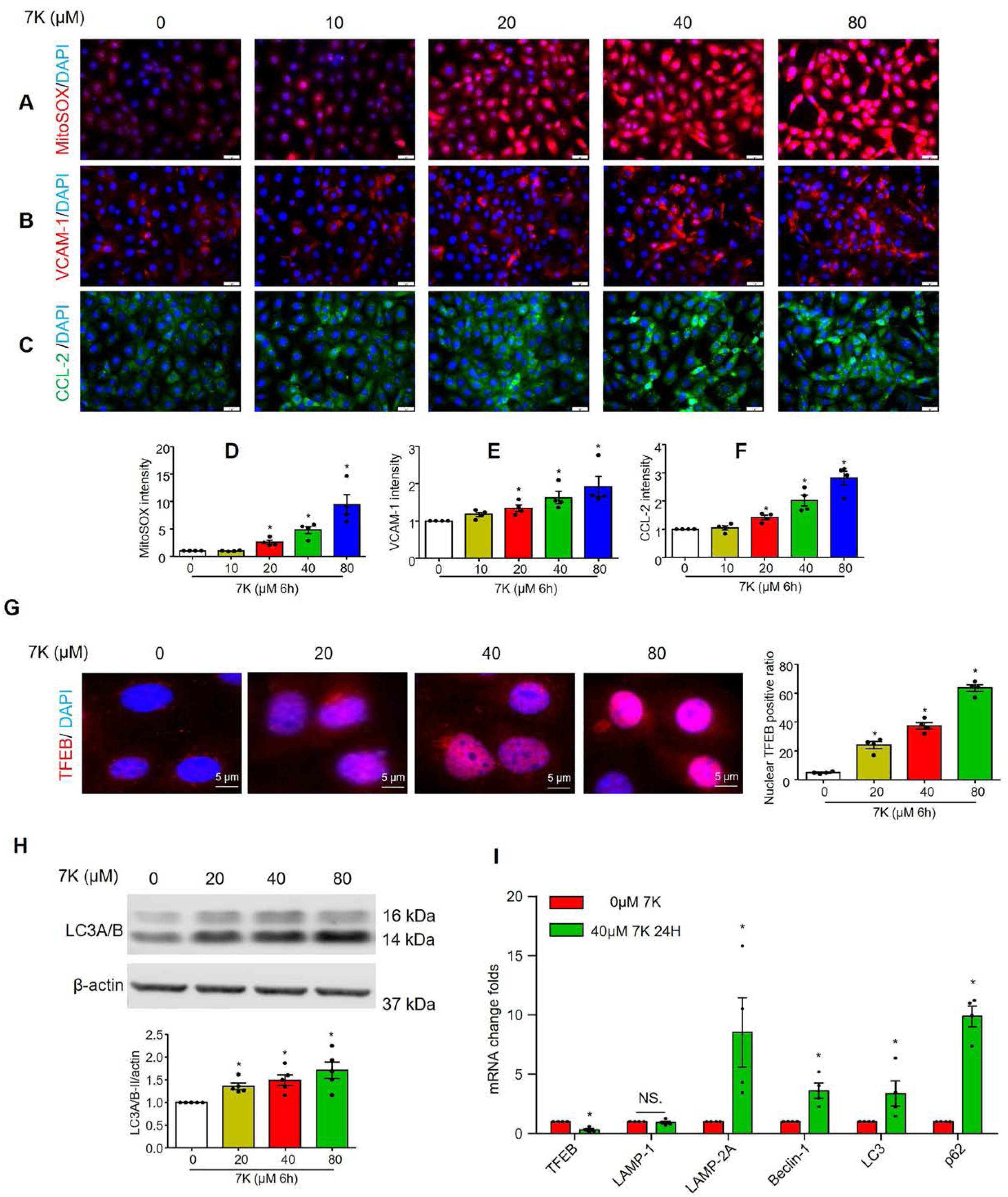
Effect of 7-ketochlesterol on mitochondrial superoxide, pro-inflammatory proteins and lysosome related signaling in cultured MCECs. MCECs were cultured in low glucose DMEM with 5% FBS, and treated with 7-ketocholesterol (7K) for the indicated times. **A and D,** Representative immunofluorescence images and quantification show mitochondrial superoxide levels. Representative immunofluorescence images and quantification show the expression of pro-inflammatory proteins VCAM-1 (**B and E**) and CCL2 (**C and F**). **G,** Representative immunofluorescence images and quantification show the nuclear translocation of TFEB. Nuclei were stained with DAPI. **H,** Representative immunoblots and summarized data show the effects of 7-keto on the protein expression levels of LC3A/B-II. **I,** Real-time RT-PCR analyses of TFEB, LAMP-1, LAMP-2A, Beclin-1, LC3, and p62/SQSTM1 mRNA levels after treatment with 0 or 40µM 7-keto for 24 hours. Scale bar=20 µm. *P< 0.05 (n=4-5).

### Inhibition of lysosomal function exacerbated inflammation and cell death in cultured MCECs

To determine the role of lysosomal signaling activation in endothelial inflammation and injury, we inhibited lysosomal function in MCECs using bafilomycin A1, a selective inhibitor of vacuolar H+ ATPases (V-ATPases) on the lysosomal membrane. Bafilomycin A1 inhibits V-ATPases and leads to the impaired acidification of lysosomes ^35^. Impaired lysosomal acidification may result in the inhibition of TRPML1, an upstream event of TFEB activation ^36–38^. It was found that bafilomycin A1 significantly inhibited 7K-induced TFEB nuclear translocation **(Fig.9A)**. We then examined the inflammatory and injury responses in MCECs by staining cells with VCAM-1, FLICA (a green fluorescent probe for activated caspase-1) ^39–41^, or propidium iodide (PI) (for detecting dead cells). We observed that bafilomycin A1 enhanced the 7K-induced increase in VCAM-1 expression (**Fig.9B**), FLICA+ casapase-1 activity (**Fig.9C and 9D**), and PI+ dead cells (**Fig.9C and 9E**). We also demonstrated that 7K increased the number of pyroptotic (FLICA+/PI+) (**Fig.9F**) and non-pyroptotic (FLICA-/PI+) (**Fig.9G**) dead cells, while bafilomycin A1 further increased the number of dead cells. By assaying cell viability, we confirmed that bafilomycin A1 exacerbated 7K-induced cell death **(Fig.9H)**. Taken together, these results suggest that TFEB-mediated lysosomal signaling protects against 7K-induced inflammatory responses and injury in cultured MCECs.

**Figure 9.**
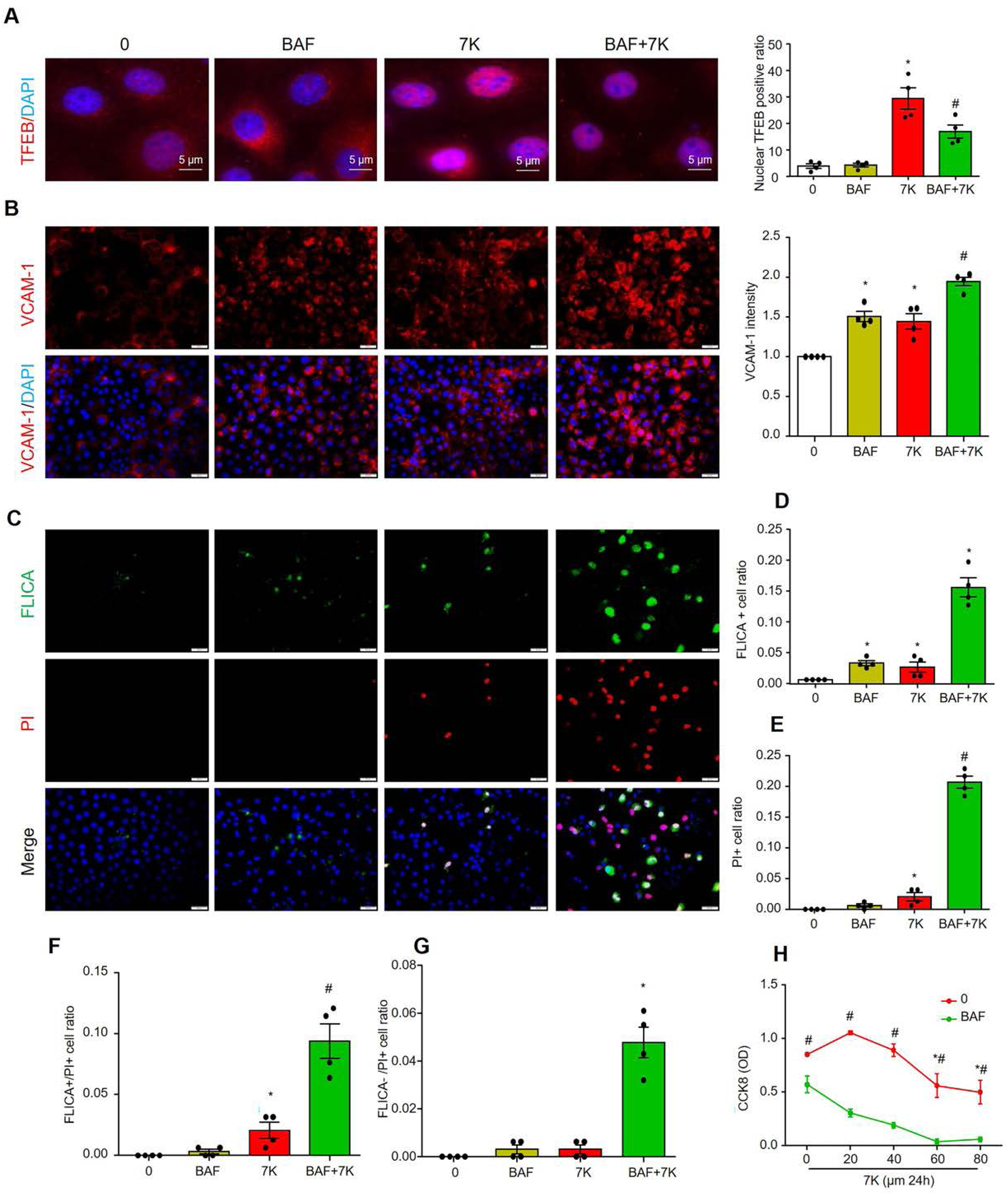
Inhibition of lysosome signaling by bafilomycin A1 exacerbates 7K-induced inflammation and cell death in cultured MCECs. MCECs were cultured and treated in low glucose DMEM with 5% FBS, pretreated with or without 50 nM bafilomycin A1 (BAF) for 1 hour, and then co-treated with or without 40µM 7-keto for 6 hours. **A,** Representative immunofluorescence images and quantification show the nuclear translocation of TFEB. Nuclei were stained with DAPI. **B-G,** MCECs are treated in low glucose DMEM with 1% FBS for 2 hours before pretreatment with or without 50 nM BAF for 1 hour, and then the cells are co-treated with or without 40 uM 7-keto for 24 hours. **B**, representative images of VCAM-1 and summarized data. **C-G**, representative images of FLICA/PI staining and summarized data. **H**, cell numbers were detected by using CCK8 kit. Scale bar=20 µm. * vs 0, # vs BAF or 7-keto, *P*< 0.05 (n=4).

### Ezetimibe activated TFEB and attenuated mitochondrial ROS and inflammation in cultured MCECs

Finally, we investigated whether ezetimibe could directly affect ECs in addition to its cholesterol-lowing effects. We found that ezetimibe dose-dependently increased TFEB nuclear translocation (**Fig.10A**) and further enhanced 7K-induced TFEB nuclear translocation (**Fig.10B**). Moreover, ezetimibe attenuated 7K-induced mitochondrial ROS production (**Fig.10C**), as well as the expression of VCAM-1 (**Fig.10D**) and CCL-2 (**Fig.10E**), and monocyte adhesion (**Fig.10F**). These results suggest that ezetimibe acts synergistically with 7K to increase TFEB-mediated lysosomal signaling and prevent 7K-induced mitochondrial ROS and inflammation in cultured MCECs.

**Figure 10.**
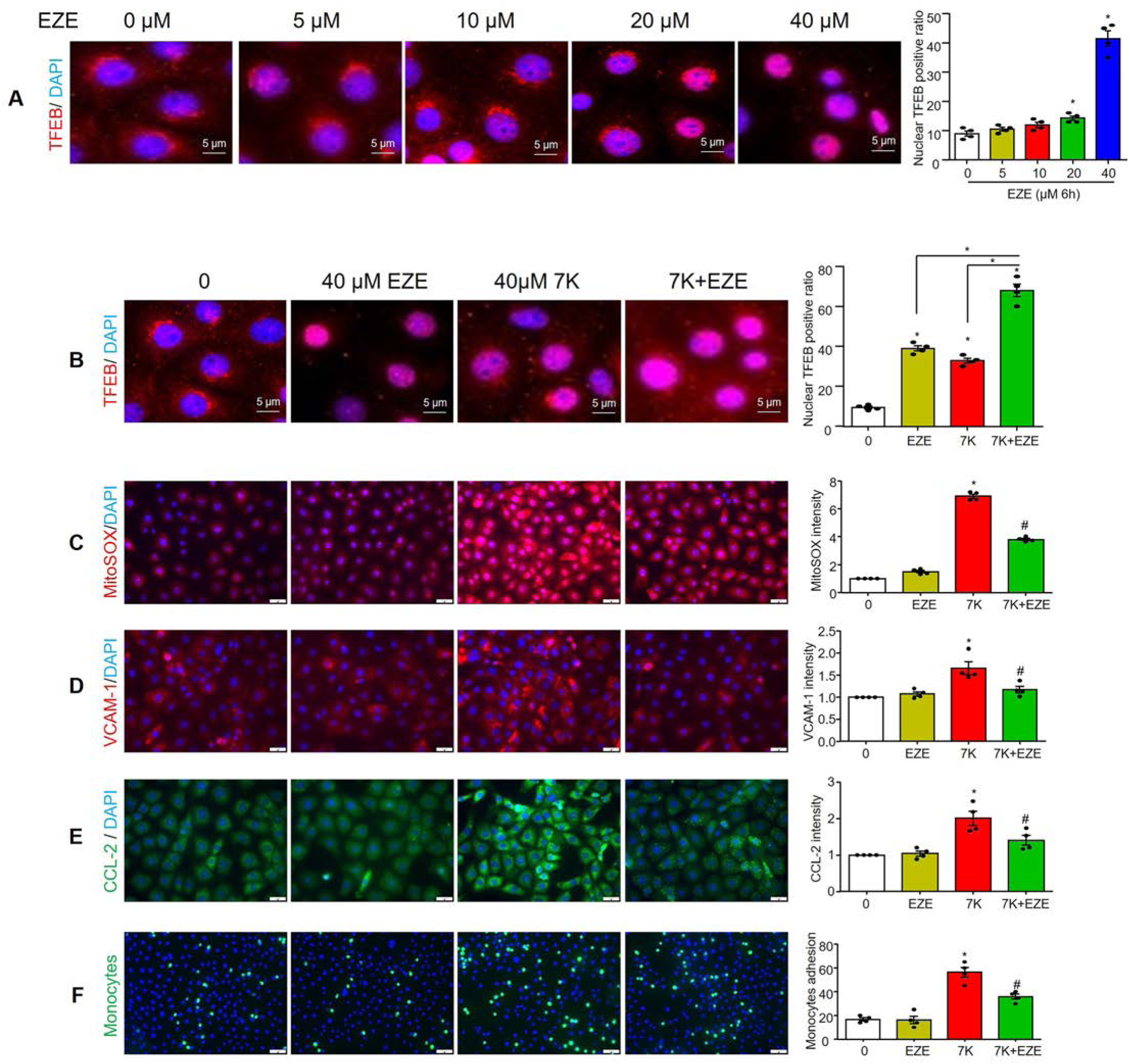
Ezetimibe attenuated 7K-induced mitochondrial superoxide, inflammation, and monocyte adhesion via increasing TFEB nuclear translocation in cultured MCECs. MCECs were cultured in low glucose DMEM with 5% FBS, then treated with ezetimibe (EZE) with or without 7K for the indicated time. **A,** Representative immunofluorescence images and quantification show the effect of ezetimibe on TFEB nuclear translocation. **B,** Representative immunofluorescence images and quantification show the effect of ezetimibe and 7-keto on TFEB nuclear translocation. Representative immunofluorescence images and quantification of mitochondrial superoxide (**C**), pro-inflammatory proteins VCAM-1 (**D**) and CCL2 (**E**), and monocyte adhesion (**F**). Scale bar=20 µm. * vs 0, # vs 7-keto, *P*< 0.05 (n=4).

## Discussion

An increasing number of studies have shown that the prevalence of CMD is high in patients diagnosed with hypercholesterolemia ^29,42–46^. Even in hypercholesterolemia patients without obstructive CAD or heart failure, CMD with reduced CFR has been observed ^42–44,46^. Despite the clinical significance of this disease, current management guidelines in the United States do not provide specific recommendations for the treatment of CMD caused by hypercholesterolemia ^5,45^. One reason for this is that preclinical studies of hypercholesterolemia and CMD are significantly less extensive than those of obstructive vascular diseases such as atherosclerosis. A recent study reported that hypercholesterolemia resulted in reduced endomyocardial CFR and capillary density in a mini-pig model without coronary stenosis ^47,48^. Moreover, a high-cholesterol diet has been shown to induce advanced occlusive coronary atherothrombosis and heart failure, leading to increased mortality, in scavenger receptor class B type 1 (SR-B1) and low-density lipoprotein receptor (LDLR) double knockout mice ^49,50^. These findings have established a clear link between hypercholesterolemia and CMD in animal studies and their contribution to obstructive CAD ^49–54^. However, the precise pathophysiological mechanisms underlying the initiation and progression of CMD caused by hypercholesterolemia remain unclear and understudied. In the present study, we established a hypercholesterolemia model in mice by feeding them a high-fat and high-cholesterol Paigen’s diet (PD). Non-invasive imaging technique showed that PD decreased hyperemic coronary blood flow velocity and CFR as time increased (**Fig.1**). CFR began to decrease after 6-8 weeks PD treatment, but no changes in cardiac remodeling or function were observed. Therefore, we used this model throughout this study to explore the early effects of hypercholesterolemia on the coronary microcirculation and myocardial tissue before the onset of cardiac remodeling and dysfunction.

Next, we characterized PD-induced CMD by evaluating a range of cardiovascular-related pathological changes in the coronary arterioles, capillaries, or myocardium, including arterial wall thickening and lipid accumulation, microvascular rarefaction, thrombosis, and inflammation. Histopathological analysis revealed that PD-induced CMD was not associated with coronary arterioles wall remodeling that may cause vascular lumen reduction and insufficient blood perfusion (**Fig.2A**). Moreover, capillary density and pericyte coverage were not affected by PD (**Fig.2B**), suggesting that PD was insufficient to induce microvascular rarefaction and decrease blood perfusion. These data suggest that PD-induced CMD may result from functional, rather than structural, changes in the coronary microcirculation.

Endothelial dysfunction is associated with increased permeability of plasma lipoproteins to the endothelium and their retention in the subendothelial area ^55,56^. Increased retention of lipoproteins in the subendothelial area may lead to the uptake of lipoproteins by resident macrophages or nearby ECs or SMCs ^55,56^. The lipid content of the uptaken lipoproteins is then stored intracellularly as lipid droplets (LDs) ^56,57^. These mechanisms may lead to lipid deposition in the vascular wall, forming fatty streaks at an early stage of occlusive vascular diseases ^55,56,58^. Since hypercholesterolemia is a common risk factor for CMD and occlusive CAD, we analyzed lipid deposition in coronary arterioles by Oil Red O staining. The results showed that the incidence of fatty streak formation in the coronary arterioles wall was increased, with fatty streaked arterioles observed in 3 out of 8 mice examined (**Fig.2C**). Gene expression analysis further showed that PD treatment tends to increase lipogenesis and LD formation in the vessel wall (**Fig.3A**). The specific location of the lipid deposition is unclear, but it may accumulate in the subendothelium or in vascular wall cells, including macrophages, ECs, or SMCs. In particular, recent studies have shown that LDs in ECs promote atherosclerosis and hypertension in mice ^59,60^. Therefore, it is possible that PD increases LD accumulation in coronary arterioles ECs, contributing to endothelial dysfunction and CMD, which deserves further investigation. Nonetheless, to our knowledge, these data for the first time demonstrates that PD-induced CMD is associated with increased lipid deposition in the vessel wall of coronary arterioles in mice.

Recently, studies have reported that PD induced severe coronary atherothrombosis in SR-B1 and LDLR double knockout mice ^49,50^. These knockout mice have plasma cholesterol levels of approximately 750 mg/dl, an extremely high concentration not usually observed in patients with metabolic disorders ^61^. In contrast, we and others have recently reported that wild-type mice fed with PD developed a mild hypercholesterolemia as early as 2 weeks, with a two-fold increase in plasma cholesterol levels compared to ND-fed controls ^34,62^, which may better represent early-stage metabolic disorders. Interestingly, the present study showed that CD41-positive platelet aggregates were found in the lumen of coronary arterioles in 2 out of 8 mice fed with PD, but not in ND-fed controls (**Fig.2D**). The increased incidence of platelet aggregates was consistent with the upregulated gene expression of tissue factor in the heart or aortic tissues (**Fig.3B**). Therefore, these results suggest that mild hypercholesterolemia induced by PD promotes CMD, accompanied by a shift in coronary arterioles ECs from an anti-thrombotic to pro-thrombotic state.

Accumulating evidence suggests that the activation of the endothelial NLRP3 inflammasome serves as a critical initiating mechanism in the pathogenesis of vasculopathies associated with metabolic disorders, including hypercholesterolemia, hyperglycemia, and dyslipidemia, as well as other conditions such as smoking and sepsis ^30,40,63–65^. Endothelial inflammasome activation leads to increased activity of caspase-1, an inflammatory caspase, and production of pro-inflammatory cytokines such as IL-1β and IL-18. In addition, endothelial inflammasome activation may lead to caspase-11 activation and GSDMD-dependent pore formation in the plasma membrane, causing pyroptotic cell death. HMGB1 is being increasingly recognized as a key DAMP molecule actively released by immune cells or passively released from dying cells upon infection or sterile inflammation ^66^. Our recent studies have shown that endothelial NLRP3 inflammasome activation promotes HMGB1 release from ECs in response to various cardiovascular risk factors ^40,41,63,64,67–69^. HMGB1 can activate RAGE or TLR2/4 receptors to initiate various inflammatory responses or cell damage ^70^. In particular, HMGB1 has been shown to be a potent vascular permeability factor that can lead to the loss of endothelial barrier function during metabolic disorders^71,72^. Upregulation of adhesion molecules, including VCAM-1 and ICAM-1, represents the initial inflammatory response of the ECs, which mediate the adhesion of leukocytes (e.g. lymphocytes, monocytes, eosinophils, and basophils) to the vascular endothelium ^73^. Recent studies have revealed that IL1β, IL-18, and HMGB1 can upregulate the expression of these adhesion molecules in ECs ^74,75^. PD has been shown to increase systemic inflammation in mice ^76^, but its effect on inflammation in CMD has not been investigated. In the present study, we found that PD significantly increased the activation of the endothelial inflammasome and inflammatory responses in coronary arterioles, as indicated by elevated caspase-1, HMGB1, and VCAM-1 (**Fig.2E-G**). In addition, there were more CD45-positive immune cells in the myocardium of PD-fed mice than ND-fed controls (**Fig.2H**), suggesting that increased leukocyte adhesion and transendothelial migration contribute to cardiac inflammation. This increase in cardiac inflammation was also confirmed by our gene expression analysis of various inflammatory mediators such as IL-6, IL-8, and TNF-α (**Fig.3C**). Interestingly, recent studies have shown that inflammatory cytokines such as TNFα promote LD formation in ECs ^77,78^. In addition, these inflammatory cytokines increase the expression of chemokines, adhesion molecules, and thrombotic modulators such as tissue factor, which may also facilitate interactions with platelets and their activation and aggregation ^17,79–81^. In this regard, the activation of the endothelial inflammasome and inflammatory responses may crosstalk with coronary lipid accumulation and thrombosis. These findings support the view that hypercholesterolemia upregulates the activation of the endothelial NLRP3 inflammasome in the coronary microcirculation and promotes the development of CMD, and that cardiac inflammation may be an early effect associated with CMD.

Another important finding of the present study is that PD-induced CMD is associated with the upregulation of lysosomal signaling pathways. Specifically, we observed upregulation of lysosomal pathway proteins, including TFEB, LAMP1, LAMP2A, and TRPML1, in the endothelium of coronary arterioles (**Fig.4**). Lysosomal signaling pathways play a critical role in regulating endothelial homeostasis under various metabolic stresses ^82,83^. For example, lysosomal function is integral to cholesterol homeostasis and is a key processing center for LDL-derived cholesterol ^84^. Within lysosomes, cholesterol esters are hydrolyzed by lysosomal acid lipase (LAL), and free cholesterols interact with NPC1/2, and are transported to extra-lysosomal destinations ^85^. Lysosomal membrane proteins LAMP1 and LAMP2A are essential for lysosomal function and LAMP-1/2 deficiency causes embryonic lethality in mice, accompanied by the accumulation of cholesterol and autophagic vacuoles in various tissues, including ECs and Schwann cells ^86^. Overexpression of LAMP-2A, but not LAMP-1, rescues cholesterol accumulation in LAMP-1/2 double-deficient fibroblasts ^86^. In addition, genetic deletion of LAMP-2 leads to luminal stenosis and medial thickening in coronary arteries ^87^. In fact, LAMP-2A-dependent chaperone-mediated autophagy plays a critical role in lipophagy, a lysosome-dependent process of LD catabolism ^88^. These previous findings highlight the more significant role of LAMP-2 in lysosomal function and lipid handling compared to LAMP-1. Furthermore, TRPML1 is a lysosomal Ca^2+^ channel, and mutations in TRPML1 cause mucolipidosis type IV (MLIV), a severe lysosomal storage disorder ^89^. Increased TRPML1 activity promotes lysosomal trafficking and fusion with intracellular organelles, including multivesicular bodies or autophagosomes, thereby facilitating lysosome-dependent degradation of these organelles and their lipid and protein contents ^90^. TRPML1-mediated lysosomal Ca^2+^ release also activates calcineurin, which in turn dephosphorylates TFEB, leading to its nuclear translocation ^91^. TFEB is a master transcription factor that controls genes involved in lysosomal and autophagy pathways. TFEB has been shown to be involved in angiogenesis and EC survival ^92^. We recently demonstrated that simvastatin induces TFEB-mediated lysosomal biogenesis, thereby preventing lipotoxicity-induced NLRP3 inflammasome activation and injury in ECs ^83^. In addition to ECs, TFEB suppression also contributes to SMC dedifferentiation, whereas activation of TFEB-lysosomal signaling promotes SMC differentiation and inhibits neointima formation under hypercholesterolemia ^31,34^. Therefore, based on these previous observations, we speculate that hypercholesterolemia upregulates lysosome signaling as an early adaptive response to lipotoxicity, and alleviates lipotoxicity-induced cellular stresses, including ER stress and mitochondrial stresses, thereby contributing to the maintenance of coronary microcirculatory homeostasis.

To date, there are no established therapeutic interventions that have been clearly proven to be effective in treating CMD. Current treatments are mainly aimed at alleviating symptoms and reducing the risk of adverse events. Commonly used drugs include aspirin, statins, angiotensin-converting enzyme inhibitors (ACEi), and angiotensin receptor blockers (ARBs) ^5^. In addition, ezetimibe has been shown to improve atherogenic lipid profiles, insulin resistance, and liver dysfunction in patients with hypercholesterolemia ^93–97^. In terms of its effect on CAD, ezetimibe, either alone or in combination with simvastatin, has demonstrated the potential to improve endothelial and microvascular function in patients with glycemic disorders and CAD ^98–100^. Animal studies have also shown that ezetimibe reduces atherosclerosis and vascular inflammation ^101–104^. Notably, ezetimibe showed a more pronounced inhibitory effect on total oxysterols, 7-ketocholesterol, and 27-hydroxycholesterol than on total cholesterol in mice with hypercholesterolemia ^100^. In the present study, we demonstrated that ezetimibe administration significantly alleviated PD-induced steatohepatitis and hypercholesterolemia (**Fig.5**), and prevented PD-induced CMD and inflammatory responses in the coronary microcirculation (**Fig.6 and Fig.7**). Our results provide the first evidence that ezetimibe may be a promising candidate for the treatment and prevention of hypercholesterolemia-induced CMD.

Next, we further elucidated the functional significance of upregulated TFEB-lysosomal signaling in endothelial inflammation and injury in the coronary microcirculation *in vitro*. A 7-ketocholesterol (7K)-induced cell injury model was established in cultured MCECs. 7K is the major oxidation product of cholesterol found in human atherosclerotic plaques and has shown greater atherogenic potential than cholesterol in animal studies ^105^. Previous reports have revealed that 7K induces mitochondrial dysfunction in ECs, characterized by altered mitochondria morphology, decreased mitochondrial membrane potential, and elevated mitochondrial superoxide levels ^106–109^. In addition, 7K has been shown to have significant pro-inflammatory effects both *in vitro* and *in vivo* ^110–113^. We have also recently demonstrated that 7K causes endothelial dysfunction and injury associated with increased oxidative stress, NLRP3 inflammasome activation, and inflammation in cultured murine ECs isolated from large carotid arteries ^39,114^. The function and integrity of lysosomes are essential for maintaining cellular homeostasis ^115^, and lysosomal dysfunction and damage can trigger or promote inflammasome activation, ER stress, or mitochondrial injury in ECs under various risk factors, including free fatty acids, cholesterol crystals, and lactobacillus casei wall components ^64,68,69,116–120^. In this study, we demonstrated for the first time that 7K induced mitochondrial ROS production, inflammation, and cellular death in cultured MCECs (**Fig.8**). Moreover, consistent with our animal studies, we found that 7K also upregulated TFEB-mediated lysosome signaling (**Fig.8**). Importantly, inhibition of lysosomal function with bafilomycin A1 suppressed TFEB activation and exacerbated 7K-induced inflammation and cell death (**Fig.9**). Therefore, our results support the view that upregulated TFEB-mediated lysosomal signaling in ECs serves as an adaptive protective mechanism to mitigate the deleterious effects associated with lipotoxicity, including mitochondrial oxidative stress, inflammatory responses, and cellular damage. The present study did not attempt to decipher the underlying mechanisms of activation of TFEB-lysosomal signaling under metabolic stress or lipotoxic conditions. Nuclear translocation of TFEB requires its dephosphorylation by inhibition of protein kinases, such as mTORC1, or activation of protein phosphatases such as calcineurin and protein phosphatase 2A (PP2A) ^121–123^. Interestingly, sustained elevated ROS levels have been shown to inhibit mTORC1 and enhance calcineurin or PP2A activities ^121,124^. We recently demonstrated that 7K triggers TRPML1-mediated lysosome-Ca^2+^ release in SMCs ^125^. The lysosomal Ca^2+^ release may activate calcineurin and in turn promote TFEB activation ^125,126^. Therefore, intracellular ROS levels and TRPML1-Ca^2+^ release may be involved in TFEB activation in MCECs, which deserves further investigation in future studies.

Lastly, another important finding of this study is that ezetimibe could synergize with 7K to increase TFEB-lysosomal signaling and prevent 7K-induced mitochondrial ROS and inflammation in cultured MCECs (**Fig.10**). Ezetimibe is a pharmacological inhibitor of NPC1L1, a cholesterol transporter that is highly expressed in the apical membrane of enterocytes and the canalicular membrane of hepatocytes. The main mechanism of ezetimibe’s beneficial effects is inhibiting intestinal and hepatic cholesterol absorption through NPC1L1, thereby lowering cholesterol ^127^. Interestingly, recent studies have shown that ezetimibe could inhibit oxidative stress, urokinase-type plasminogen activator receptor (uPAR) expression, or IL-6 secretion in human umbilical vein endothelial cells (HUVECs) ^128,129^, and have direct vasodilatory effects on rat mesenteric resistance arteries ^130^. These previous studies and the findings of this study support the view that ezetimibe can affect the cardiovascular system independently of its lipid-lowering effects, and that upregulation of TFEB-lysosomal signaling may represent one of the non-lipid-lowering effects of ezetimibe. As NPC1L1 is highly expressed in the liver and small intestine but relatively low in other sites, the mechanism of action of ezetimibe in the cardiovascular system may involve targets other than NPC1L1. Ezetimibe has recently been shown to activate TFEB in hepatocytes via stimulation of AMPK ^131^. The exact mechanism by which ezetimibe activates TFEB-lysosomal signaling and the possible role of AMPK in this effect warrants further investigation.

In summary, the present study demonstrated that CMD develops in a mouse model of mild hypercholesterolemia before the onset of cardiac remodeling and dysfunction. The most notable pathological changes associated with CMD were enhanced inflammation in the coronary microcirculation and myocardium. Moreover, the progression of CMD was accompanied by adaptive upregulation of TFEB-lysosomal signaling in coronary ECs, which may attenuate the inflammation and injury in ECs. Ezetimibe, an FDA-approved cholesterol-lowering drug, effectively reversed these deleterious effects, which is thought to be achieved by activating TFEB-lysosomal signaling to exert its non-lipid-lowering effect. Targeting TFEB-lysosomal signaling may be a promising therapeutic strategy to prevent hypercholesterolemia-induced CMD.

## Acknowledgment

This work was supported by the grants from the National Institutes of Health (R01HL150007, R01HL122937).

## Abbreviations

CMD: Coronary microvascular dysfunction
PD: Paigen’s diet
ND: Normal diet
7K: 7-ketocholesterol
MCECs: Mouse cardiac endothelial cells
BAF: Bafilomycin A1
SR-B1: Scavenger receptor class B type 1
LDLR: Low-density lipoprotein receptor
TFEB: Transcriptional factor EB
LAMP-1: Lysosomal-associated membrane protein 1
LAMP-2A: Lysosome-associated membrane protein type 2A
LC3A/B: Microtubule-associated proteins 1A/1B light chain 3A/B
p62: p62/SQSTM1
TRPML-1: Transient receptor potential cation channel, mucolipin subfamily, member 1
HMGB-1: High mobility group box 1
VCAM-1: Vascular cell adhesion molecule 1
ISO: Isoflurane
CFR: Coronary flow reserve
LVAW: Left ventricular anterior wall
LVID: Left ventricular internal end
LVPW: Left ventricular posterior wall
LVEF: Left ventricular ejection fraction
LVFS: Left ventricular shortening fraction
GPAT-4: Glycerol-3-phosphate acyltransferase 4
AGPAT-2: 1-Acylglycerol-3-Phosphate O-acyltransferase 2
DGAT-1: Diacylglycerol O-acyltransferase 1
DGAT-2: Diacylglycerol O-acyltransferase 2
ACAT-1: Acetyl-CoA acetyltransferase 1
ACAT-2: Acetyl-CoA acetyltransferase 2
TFPI: Tissue factor pathway inhibitor
PAI-1: Plasminogen activator inhibitor 1
PAR-1: Protease activated receptor 1
NLRP3: NLR family pyrin domain containing 3
NLRP1A: NLR family pyrin domain containing 1A
ASC: PYD and CARD domain containing
ICAM-1: Intercellular adhesion molecule 1
TNF: Tumor necrosis factor
GSDMD: Gasdermin D
SMPD1: Sphingomyelin phosphodiesterase 1
CCL2: C-C motif chemokine ligand 2

